# Specific splice junction detection in single cells with SICILIAN

**DOI:** 10.1101/2020.04.14.041905

**Authors:** Roozbeh Dehghannasiri, Julia Eve Olivieri, Julia Salzman

## Abstract

Precise splice junction calls are currently unavailable in scRNA-seq pipelines such as the 10x Chromium platform but are critical for understanding single-cell biology. Here, we introduce SICILIAN, a new method that assigns statistical confidence to splice junctions from a spliced aligner to improve precision. SICILIAN’s precise splice detection achieves high accuracy on simulated data, improves concordance between matched single-cell and bulk datasets, increases agreement between biological replicates, and reliably detects un-annotated splicing in single cells, enabling the discovery of novel splicing regulation.

## Main text

Alternative splicing is essential for the specialized functions of eukaryotic cells, necessary for development ^1^, and a greater contributor to genetic disease burden than mutations ^2^. Despite the importance of splicing and massive RNA-seq data generated on single cells, the extent to which the diversity of RNA splicing in single cells is regulated and functional versus transcriptional noise remains contentious ^3^.

Given the resolution and massive number of available single-cell RNA-seq (scRNA-seq) datasets, precise quantification of splicing in single cells has great promise for discovering regulatory and functional splicing biology. While there are a number of methods developed for isoform quantification at the single-cell level ^4,5^, a problem that has not been addressed is the precise discovery of splice junctions. There is a great need for precise junction detection: spliced aligners are designed for bulk RNA-seq and, in addition, generate many artifacts ^6–10^, which will be referred to in this paper as “false positives”. The problem is exacerbated in scRNA-seq analysis due to the high-level and single-cell-specific biochemical noise and multiple testing errors arising from the analysis of thousands of cells. This problem is typically addressed through the use of simple filters on junction calls, which remove many true positives, especially when the data is sparse such as scRNA-seq. Because of these challenges, there is debate regarding whether 10x Chromium (10x) could be used for reliable de novo splice junction detection ^11,12^, despite the presence of a large number of junctional reads in the datasets generated by 10x protocol (Figure 2A). Overall, existing approaches to splicing analysis in the scRNA-seq data either lack sufficient sensitivity to identify splice junctions or specificity to identify false positives.

**Figure 1:**
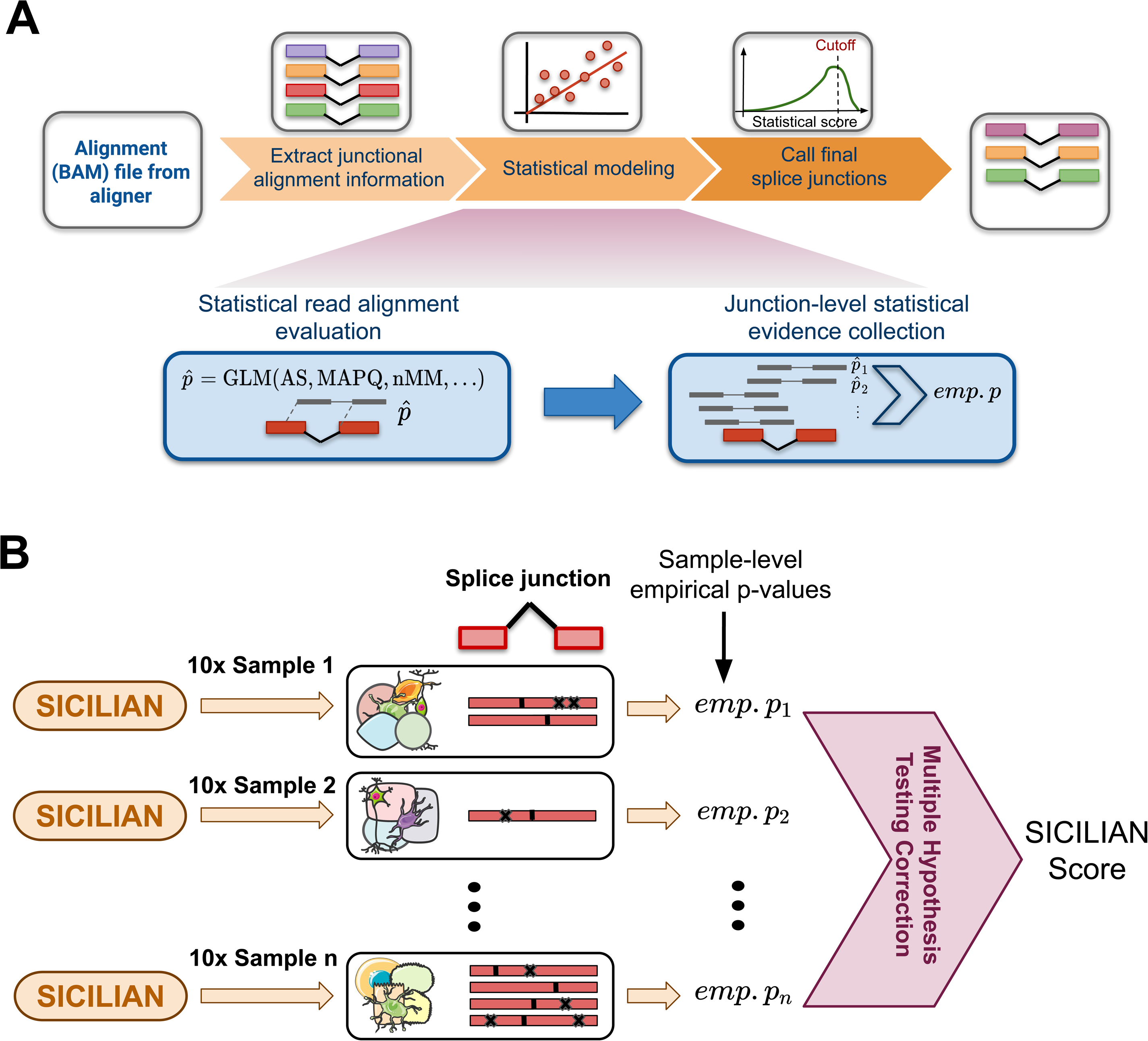
Overview of the SICILIAN statistical framework. (A) SICILIAN takes the alignment information file (usually in the form of a BAM file) from a spliced aligner such as STAR and then deploys its statistical modeling to assign a statistical score to each junction. (B) SICILIAN utilizes the cell-level statistical scores (empirical p-values) for each junction across 10x samples to correct for increased false discovery rates due to multiple hypothesis testing. The corrected score is called the “SICILIAN score” and can be used to consistently call junctions across cells.

**Figure 2:**
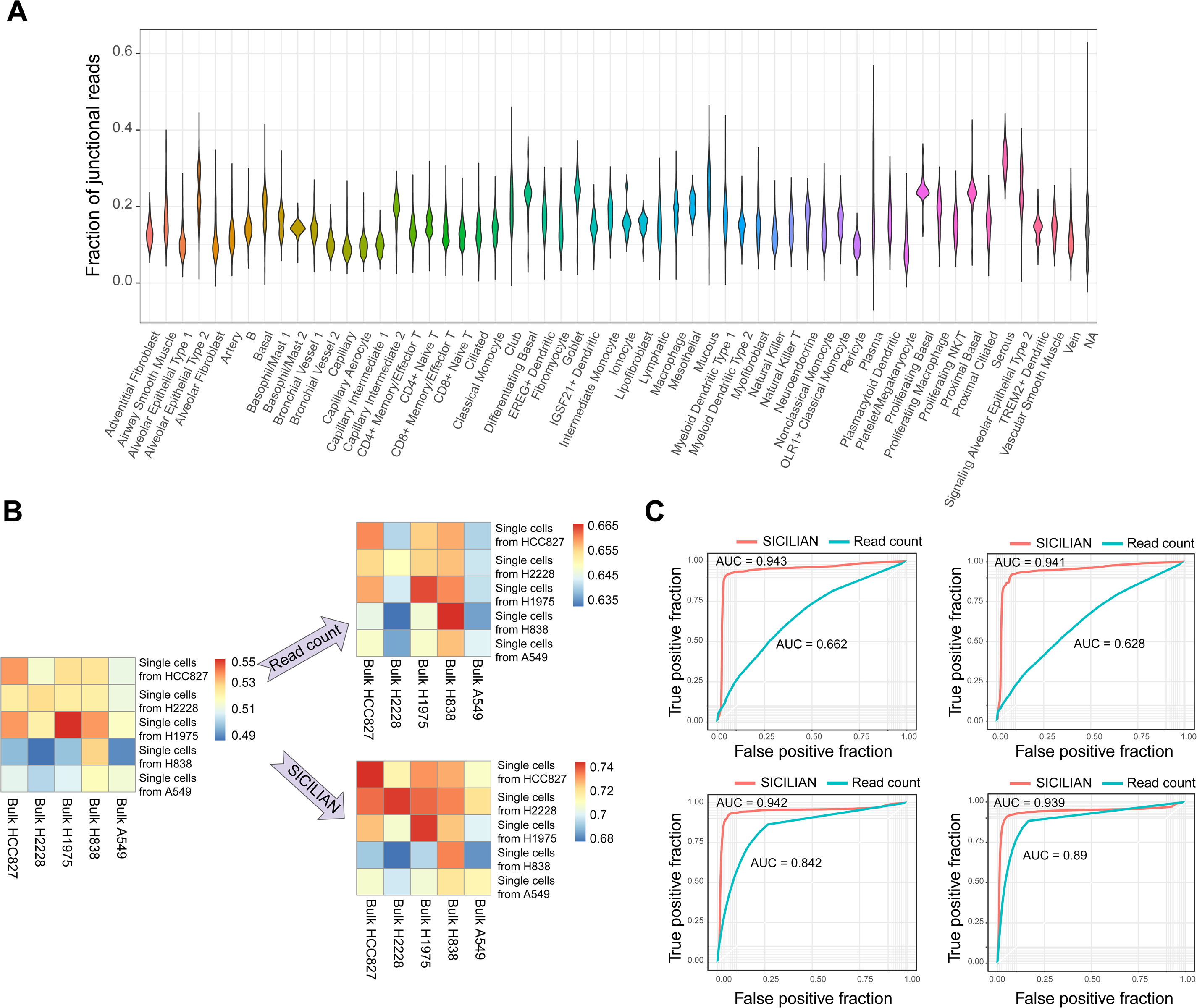
SICILIAN performance evaluation using benchmarking datasets. (A) High and variable fraction of junctional reads across diverse cell types in the HLCA dataset ^16^. Each violin plot shows the fraction of mapped reads in each cell (within a cell type) that are junctional. (B) SICILIAN improves the concordance between detected splicing junctions in single cells and bulk cell lines. (C) ROC curves by SICILIAN and read count criteria for four simulated datasets ^9,18^ (the top two based on data from ^9^ and the bottom two based on data from ^18^).

In this paper, we introduce SICILIAN (SIngle Cell precIse spLice estImAtioN), a statistical wrapper for precise splice junction quantification in single cells. SICILIAN deconvolves biochemical noise, generated during library preparation, and computational noise, generated by the spliced aligner while mapping reads to the genome, which are both highly prevalent in scRNA-seq and can lead to false-positive junctions. To identify spliced alignments that are erroneously reported by the aligner due to this combined noise, SICILIAN employs generalized linear statistical modeling, with predictors being read mapping features. In this paper, we use STAR ^13^ as the spliced aligner in SICILIAN, though the general statistical framework can be applied to refine the splice junction calls from any spliced aligner generating a BAM file.

The SICILIAN workflow has three main steps: (1) Assign a statistical score to each junctional read’s alignment to quantify the likelihood that the read alignment is truly from RNA expression rather than artifacts; (2) Aggregate read scores to summarize the likelihood that a given junction is a true positive; (3) Report single-cell resolved junction expression quantification, corrected for multiple hypotheses testing (Methods, Figure 1A,B).

The goal of step (1) above is to statistically evaluate the confidence of the alignment for each junctional read. To do this, SICILIAN fits a penalized generalized linear model ^14^ on the input RNA-seq data, where positive and negative training classes are defined based on whether each junctional read also has a genomic alignment. Training a new model for each input dataset allows SICILIAN to adapt to batch effects. The model uses the following predictors: the number of alignments for the read, the number of bases in the longer and shorter read overhangs on each side of the junction, the alignment score adjusted by the read length, the number of mismatches, the number of soft-clipped bases, and read entropy. Read entropy is not generally appreciated as an important variable in scRNA-seq reads even though it is characteristic of technical artifacts ^15^, underlining its importance in the SICILIAN model. For example, in the 10x data from a human lung study ^16^, the average read entropy in 20% of cells is < 4 (Supplementary Figure 1A), which is much more than the fraction of low-entropy reads in bulk RNA-seq datasets, e.g., the entropy is < 4 in only 0.09% and 0.4% of reads in one simulated ^9^ and five bulk cell lines, respectively (Supplementary Figure 1B,C). In step (2), the statistical scores assigned to each junction’s aligned reads are aggregated using a Bayesian hypothesis testing framework to obtain aggregated junction-level scores. SICILIAN subsequently uses the distribution of aggregated scores for likely false positive junctions to predict an empirical p-value for each junction. Finally, in step (3), SICILIAN corrects for multiple hypotheses by taking the median of the empirical p-values for each junction across samples and reports it as the final “SICILIAN score” for the junction (Figure 1B). User-defined thresholding on this score allows for a junction to be either called or thrown out consistently across all samples. In this paper, we used a threshold of 0.15, which was selected to maximize the sum of sensitivity and specificity on the benchmarking datasets with known ground truth (Methods).

We benchmarked SICILIAN using three different types of data: matched bulk and scRNA-seq data from five human lung adenocarcinoma cell lines ^17^, simulated data with known grown truth ^9,18^, and real scRNA-seq data, where we ran SICILIAN on 36,583 lung cells from two individuals from the human lung cell atlas (HLCA) ^16^ and 16,755 lung cells from two individuals from the mouse lemur cell atlas (MLCA) studies, all sequenced using the 10x platform. We compared SICILIAN to commonly used filtering criteria in the field: all junction calls based on STAR ^13^ raw alignments, the junctions supported only by uniquely mapping reads ^19^, and calling junctions based on read counts ^20,21^.

First, we show that SICILIAN increases the concordance of junction calls on matched single-cell and bulk datasets ^17^. We define “concordance” to be the fraction of junctions detected in the single cells from each cell line that are also present in the bulk data from the same cell line. SICILIAN increases the concordance between the detected junctions from 10x and bulk RNA-seq regardless of the pairs’ cells of origin, which is consistent with SICILIAN identifying and removing scRNA-seq specific artifacts. SICILIAN improves the concordance for all cell lines (Figure 2B), e.g., for cell line HCC827, the concordance based on raw STAR calls is 0.54 and SICILIAN increases it to 0.75, while calling junctions based on a 10-read filter only increases the concordance to 0.66.

Second, SICILIAN increases prediction accuracy on four bulk simulated datasets with known ground truth ^9,18^. As there is no single-cell dataset with fully known ground truth, we resorted to bulk-level simulated datasets but ran the identical SICILIAN model on them. For these datasets, SICILIAN uniformly achieves AUCs of ∼0.94, a significant increase over the AUCs of 0.66-0.89 based on the read count criterion (Figure 2C).

Third, SICILIAN increases the proportion of annotated to unannotated junctions in all four human and mouse individuals compared to the original STAR calls. We expect an algorithm that correctly identifies false positives to enrich for annotated junctions, particularly for organisms with extensive annotation (e.g., human) ^22^. Because transcript annotations are not part of the SICILIAN model, this serves as an orthogonal measure for performance evaluation. In all four individuals, SICILIAN calls a higher proportion of annotated junctions (83.6% on average) than unannotated junctions (29.2% on average), and a higher proportion of annotated junctional reads (87.6% on average) than unannotated junctional reads (23.9% on average), excluding junctions that only appear once in the dataset (Figure 3A,B; Supplementary Figure 2). Considering annotated and unannotated junctions as surrogates for true positive and false positive junctions in human lung data, respectively, SICILIAN achieves an AUC of 0.74, while that of the read-count-based approach is 0.5 (Figure 3C).

**Figure 3:**
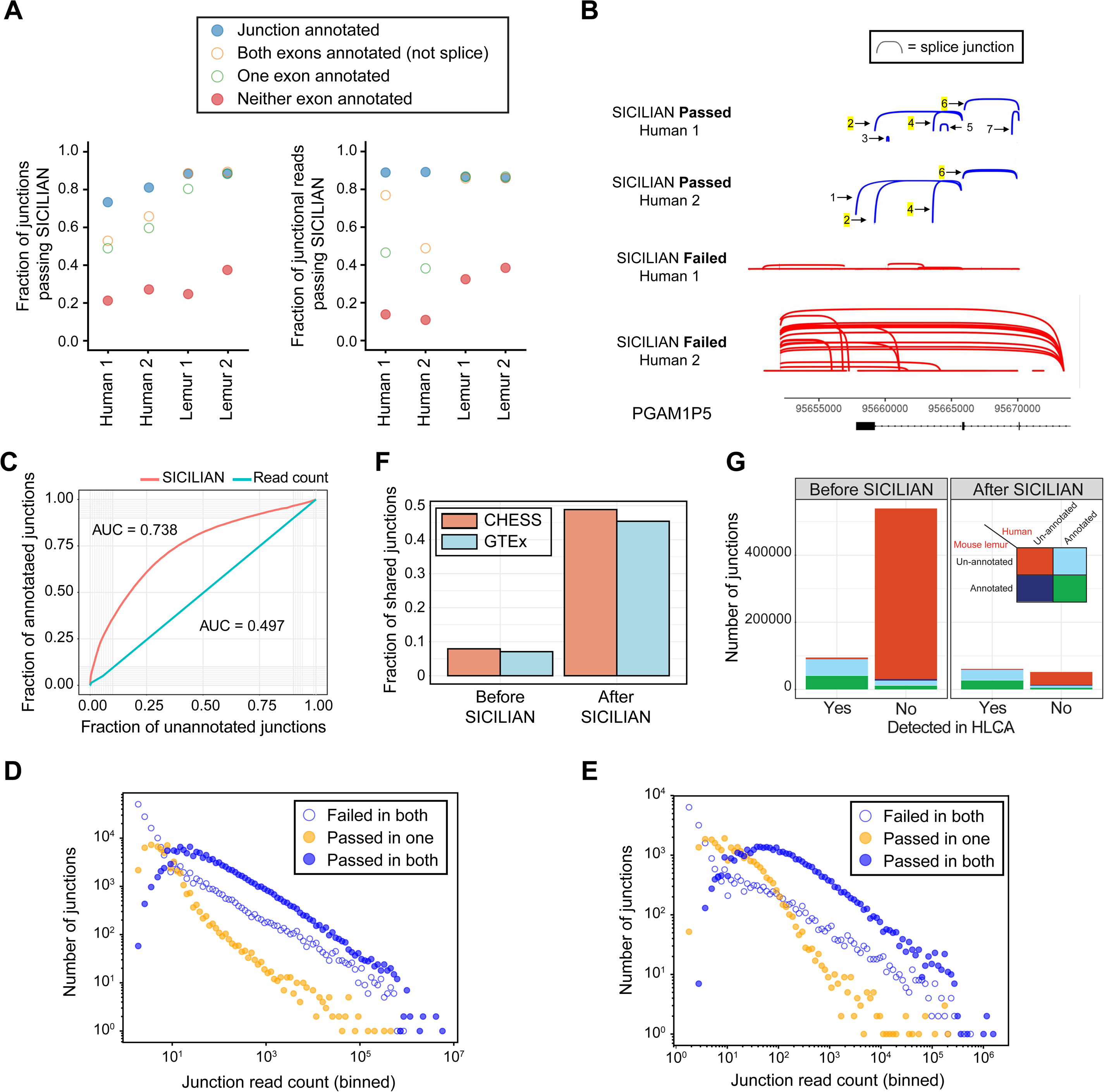
Splice junction discovery in human lung (HLCA) and mouse lemur lung (MLCA) cells. (A) SICILIAN filters out a higher proportion of unannotated junctions than annotated in all individuals from both human and mouse Lemur datasets [only junctions with at least two reads in the given dataset are plotted]. (B) Example of junctional reads identified by SICILIAN in the gene PGAM1P5 in human; annotated splicing was maintained and four new junctions were discovered (one of which was shared between individuals). (C) Better discrimination between annotated and unannotated junctions in the HLCA dataset achieved by the SICILIAN statistical criterion. (D) The number of junctions found in both human individuals that are called consistently by SICILIAN is larger than the number that are called differently in almost every case, regardless of the number of junctional reads. (E) The number of junctions called consistently between the two mouse lemur individuals is also larger than the number called inconsistently at almost every read depth after SICILIAN filtering. (F) The fraction of HLCA junctions that are found in CHESS and GTEx databases before and after applying SICILIAN to STAR raw calls. (G) The number of mouse lemur junctions orthologous junctions (found by LiftOver from Mmur3 to hg38) that have been also detected in the HLCA dataset. Junctions have been further classified based on their annotation status in the mouse lemur and human transcriptomes.

Fourth, SICILIAN increases the agreement of splicing calls between individuals. SICILIAN calls 117,684 shared junctions between individuals in HLCA, while a 10-read cutoff calls only 80,292. SICILIAN makes a consistent call (either calls or rejects the junction in both) for 83.0% of junctions. Similarly, in MLCA, SICILIAN calls 36,446 shared junctions in both individuals, while a 10-read cutoff calls only 17,798. SICILIAN consistently calls 69.9% of junctions across mouse lemur individuals. At almost all fixed junction expression levels, SICILIAN makes more consistent calls than inconsistent calls at both human and mouse lemur, emphasizing the robustness of SICILIAN across datasets (Figure 3D,E).

To further identify whether SICILIAN enriches for known junctions, we compared the junctions called by SICILIAN in HLCA with two of the most recent and precise databases of human splice junctions: CHESS ^23^ and the Genotype-Tissue Expression (GTEx) project ^24^. Only 8% and 7% of raw STAR calls are present in CHESS and GTEx, respectively, but SICILIAN increases these percentages to 48.8% and 45.4%, respectively (Figure 3F). We also looked at the junctions within each lung cell type and found that the fraction of junctions that are not present in CHESS or GTEx varies substantially across different cell types, even those with similar sequencing depth, indicating that the rate of novel splicing may vary between cell types (Supplementary Figure 3).

Finally, mouse lemur calls by SICILIAN are enriched in having annotated orthologous junctions (obtained via the LiftOver tool ^25^) in the human transcriptome, which is much more complete than the mouse lemur transcriptome ^26^ (Methods). Strikingly, more than 48.4% of un-annotated junctions called by SICILIAN are annotated in the human transcriptome, compared to only 11.2% of unfiltered STAR calls, which supports the claim that SICILIAN filtering enriches for true positive junctions. We also compare the detected junctions in MLCA and HLCA datasets: applying SICILIAN increases the fraction of junctions in mouse lemur that have been also detected in HLCA from 15% to 54% (Figure 3G).

Taken together, our results demonstrate that the SICILIAN method enables a new level of precision in spliced junction detection from single-cell platforms such as 10x. SICILIAN allows automatic junction discovery even for poorly annotated splicing programs such as rare cell types or in new model organisms. The conceptual models used in SICILIAN are also applicable to other data types such as emerging single-cell sequencing technologies and bulk and long-read sequencing.

## METHODS

### Alignment of scRNA-seq data

FASTQ files were aligned using STAR with chimSegmentMin = 10 and chimJunctionOverhangMin = 10 and the rest of the parameters were set to their defaults. Every spliced alignment (defined as a read with an “N” in its cigar string that was not chimeric) was parsed from the STAR BAM files. By collapsing spliced alignments based on their mapping positions, we obtain the “unfiltered STAR calls.” If a read had multiple spliced alignments, we only included the spliced alignment with the lowest value of the HI BAM flag to avoid double-counting reads. We also kept track of which reads had genomic alignments as needed for selecting our training datasets.

### Statistical detection of splice junctions in single-cell data

SICILIAN extracts all relevant information for the spliced alignments from the BAM file and utilizes that information to build a statistical model to distinguish truly expressed junctions from false positives due to biochemical and computational noise (Figure 1A). To build the model, SICILIAN takes advantage of the information across thousands of cells in a 10x sample to train a logistic regression model and thereby assigns a single statistical score to each extracted spliced alignment. Therefore, for each sample that SICILIAN is run on, a new model is built, allowing the model to specifically adapt to the experimental conditions of the given sequencing run. For the data analyzed in this paper, each 10x lane was modeled separately to allow the modeling of lane-specific batch effects. The statistical framework in SICILIAN can be divided into three main steps: (1) Statistical read alignment evaluation step, (2) Junction-level statistical evidence collection, and (3) Multiple hypothesis testing correction.

### Statistical read alignment evaluation

SICILIAN trains a different model for every dataset. To train the regression model, we define negative training reads as the reads that have both spliced and genomic (i.e., a contiguous mapping to a genomic region) alignments and positive training reads as the junctional reads without a genomic alignment. Our choice of negative training reads reflects the mapping profile of erroneous alignments as the vast majority of junctional reads with genomic alignments are not due to the real splice junction expression, but rather confounding factors such as genome homology and other biochemical and sequencing errors. Another advantage is that our selection criterion for training reads is independent of the predictors in the regression (because whether a spliced read also has a genomic alignment is not included as a feature in the model). Therefore, the training reads would not give too much weight to any predictor in the fitted regression model. Each positive and negative training set can have at most 10,000 junctional reads, chosen randomly among the set of reads satisfying the training reads criterion.

Let *y* be a binary variable, where *y*=1 indicates a true spliced alignment and *y*=0 indicates a false alignment. We model *p*(*y*=1), the likelihood of a true spliced alignment, with the logistic regression with penalized maximum likelihood ^14^. The predictors in the regression comprise: number of reported alignments for the read obtained from the NH tag in the BAM file (*NH*), number of mismatches obtained from the NM tag in BAM file (*nmm*), length-adjusted alignment score obtained by normalizing the alignment score from AS tag to the read length (*AS*), length of the shorter read overhang flanking the junction (*overlap*), length of the longer read overhang flanking the junction (*max_overlap*), number of the soft-clipped bases given by the S segment of the CIGAR string (*S*), and entropy of the read sequence (*entropy*):

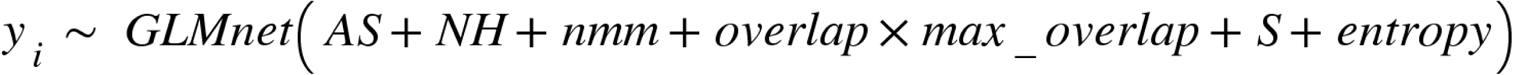

By adopting this regression model, SICILIAN evaluates spliced alignments for all cells within a 10x sample, even for those cells with low read coverage. We fitted the regression using the GLMnet R package ^14^. The fitted model is then applied to each junctional read to estimate a read-level score 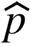, the estimated likelihood that the spliced alignment is true.

### Junction-level statistical evidence collection

The list of extracted junctions from the BAM file with their aligned reads is obtained by collapsing the spliced alignments based on their mapping position. For each junction, which corresponds to N aligned reads, the read-level scores are aggregated under a Bayesian hypothesis testing framework to obtain an aggregated junction-level score:

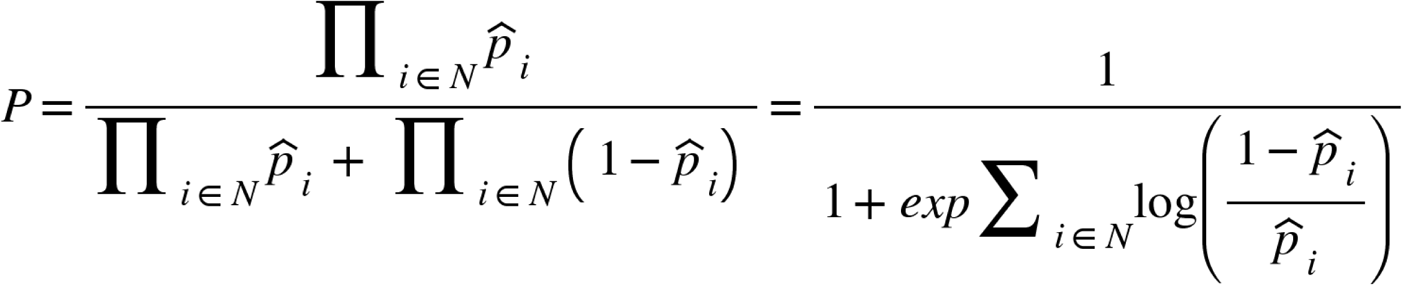

Since read-level scores are always between 0 and 1, the aggregated score would be biased against junctions with many reads even when the read alignments have high confidence. To correct this bias, for each number N of aligned reads, we build a null distribution of random aggregated scores by randomly sampling N aligned reads across all read alignments. We then compute a junction cumulative score *p*_*cum*_ for each aggregated score *P* by comparing it against the null distribution. For junctions with *N* < 15 reads, we build the empirical null distribution by computing 10,000 random aggregated scores; for junctions with *N* > 15 reads, we deploy the central limit theorem to model the null distribution as a Gaussian distribution and use it to compute the cumulative score *p*_*cum*_.

To develop a systematic approach for estimating the SICILIAN’s false discovery rate, we further computed an empirical p-value (*emp.p*) for each junction. To do so, we built a null distribution of the cumulative scores *p*_*cum*_ of the junctions in which at least 10% of the aligned reads have also a genomic alignment. Since a spliced read with a genomic alignment could be also explained by a contiguous genomic region and its reported spliced alignment is most likely due to the artifacts, we used the spliced junctions with a considerable fraction of aligned reads (at least 10%) being also genomically aligned as surrogates for false-positive junctions and used the distribution of their cumulative scores *p*_*cum*_ as the null distribution to compute an empirical p-value *emp.p* for each junction. When SICILIAN is applied to a single sample (one 10x sample), *emp.p* is the SICILIAN’s final score for each junction and junctions with *emp.p* less than a threshold (e.g. 0.1) are called by SICILIAN.

### Multiple hypothesis testing correction

When analyzing multiple 10x samples (which is typical in single-cell studies), merely using empirical p-values for detecting junctions might result in an increased false discovery rate due to multiple hypothesis testing, where each junction is tested multiple times across samples. To address this issue, SICILIAN adopts a multiple hypothesis testing correction strategy where for each junction, it collects the empirical p-values (*emp.p*) across all samples and then computes their median (or the “SICILIAN score”) as a unified criterion to decide whether the junction should be called across all 10x samples (Figure 1B). We used median because as the number of random variables increases, their median converges to their expectation; therefore, the median can be used as a consistent statistical measure that controls for multiple hypothesis tests of a junction being an artifact across samples. With this approach, if a junction possesses a significant *emp.p* in one sample just by chance but there is enough evidence in other samples that the junction is a false positive, SICILIAN would be able to correct the call by considering the junction’s *emp.p*’s in other samples, which would lead to a large median and consequently the removal of the junction. Using this system, each junction will be called consistently across all samples. For this paper, we used a cutoff of 0.15 for the SICILIAN score (or the median of *emp.p*) as this value optimized the Youden’s index (*sensitivity - specificity - 1*) on the benchmarking datasets with known ground truth.

For each sample in which junction *i* is originally present, the *emp.p* is only considered in this step if it meets several criteria: the fraction of reads with a genomic alignment that are also mapped to this junction in the sample is < 0.1, the reads mapping to the junction in this sample have different starting points for their alignment (reads aligned to a junction have different overlaps), the average length of the longest stretch of either A’s, T’s, G’s, or C’s in the reads mapping to the junction is less than 11, and the average entropy of aligned reads to the junction is greater than 3. These filters are included because any of these individual criteria provides significant evidence on its own that the junction is a false positive. If a junction “fails” any of these criteria in one sample, the *emp.p* from that sample will not be included in the set of *emp.p* used to find the median.

### Read sequence entropy

To further identify false positives by spliced aligner, the SICILIAN model also includes the entropy ^27^ for the aligned reads, a quantitative measure of how repetitive a sequence is. For example, the entropy of sequence TCACTCTCCCACACTCTCTCTCTCTCACACACACACACACACACACACACACACAC ACAC, which has many repeats of AC and TC is 2.1, while the entropy of sequence GAAAGTGTATAACTACAATCACCTAATGCCCACAAGGTACTCTGTGGATATCCCCT TGGA is 4.0. Sequence entropy is expected to be a highly informative predictor of false positive spliced alignments for two reasons: (1) reverse transcriptase or PCR enzymes are known to generate sequences of low entropy (“PCR stutter”) ^15^, and PCR crossover is common in these regions; (2) these low-entropy sequences typically map to many places in the genome ^28^. Also, the entropy could be more informative and variable in scRNA-seq compared to bulk RNA-seq (Supplementary Figure 1).

We computed the entropy for a read sequence based on the distribution of overlapping 5-mers in the sequence. For example, for ACTCCGAGTCCTCCG the list of 5-mers would be: ACTCC, CTCCG, TCCGA, CCGAG, CGAGT, GAGTC, AGTCC, GTCCT, TCCTC, CCTCC, and CTCCG. Let *k*_1_,…, *k*_*n*_ be all the 5-mers in the read sequence and C denote the set of unique 5-mers in the read, then for any *c* ∈ *C*, let be the number of times that kmer *c* appears in *k*_1_,…, *k*_*n*_ (for example, CTCCG appears twice in ACTCCGAGTCCTCCG). We define the read entropy as 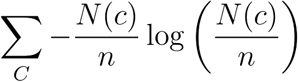.

### Mouse lemur liftover analysis

We used the UCSC LiftOver tool ^25^ (https://genome.ucsc.edu/cgi-bin/hgLiftOver) to convert the the coordinates of the junctions detected by SICILIAN in MLCA from mouse lemur (Mmur3) to human (hg38) genome assemblies and analyzed the annotation status of the orthologous junctions in the human transcriptome. We used the recommended settings for LiftOver and analyzed only those junctions that have been successfully and uniquely converted by the LiftOver tool.

### File downloads

- Human RefSeq hg38 annotation file was downloaded from: ftp://ftp.ncbi.nlm.nih.gov/refseq/H_sapiens/annotation/GRCh38_latest/refseq_identifiers/GRCh38_latest_genomic.gff.gz
- Mouse Lemur RefSeq Micmur3 assembly files were downloaded from: https://www.ncbi.nlm.nih.gov/assembly/GCF_000165445.2/
- The list of GTEx splice junctions was downloaded from GTEx Portal: https://storage.googleapis.com/gtex_analysis_v8/rna_seq_data/GTEx_Analysis_2017-06-05_v8_STARv2.5.3a_junctions.gct.gz

### Code availability

SICILIAN code is publicly available and can be accessed via a GitHuB repository: https://github.com/salzmanlab/SICILIAN. All code used for benchmarking can be found at https://github.com/salzmanlab/SICILIAN/tree/master/benchmarking.

### Data availability

The 10x benchmarking dataset ^17^ is available on SRA database (GSM3618014). The corresponding cell lines (HCC827, H1975, A549, H838, and H2228) for the benchmarking 10x dataset were downloaded from the NCI Genomic Data Commons (GDC) Legacy Archive (https://portal.gdc.cancer.gov/legacy-archive). The simulated benchmarking datasets ^9^ were downloaded from ArrayExpress (accession number: E-MTAB-1728).The HISAT simulated datasets ^18^ were downloaded from http://www.ccb.jhu.edu/software/hisat/downloads/hisat-suppl/reads_perfect.tar.gz and http://www.ccb.jhu.edu/software/hisat/downloads/hisat-suppl/reads_mismatch.tar.gz. The human lung scRNA-seq data used here was generated through the Human Lung Cell Atlas. ^29^ The mouse lemur single-cell RNA-seq data used in this study was generated as part of the Tabula Microcebus consortium.

## Acknowledgments

The authors thank Mark Krasnow and Stephen Quake for many useful discussions throughout the project; Kyle Travaglini and Camille Erzan for useful discussions and providing access to the sequencing data; Rob Bierman, Elisabeth Meyer and Kyle Travaglini for critical reading of the manuscript; the Salzman Lab for many useful discussions throughout the project. J.O. is supported by the National Science Foundation Graduate Research Fellowship under Grant No. DGE-1656518 and a Stanford Graduate Fellowship. R.D. is supported by Cancer Systems Biology Scholars Program Grant R25 CA180993. J.S. is supported by the National Institute of General Medical Sciences Grant R01 GM116847, NSF Faculty Early Career Development Program Award MCB1552196, a McCormick–Gabilan Fellowship, and a Baxter Family Fellowship. J.S. is also an Alfred P. Sloan Fellow in Computational & Evolutionary Molecular Biology.

## List of Figures

**Supplementary Figure 1.**
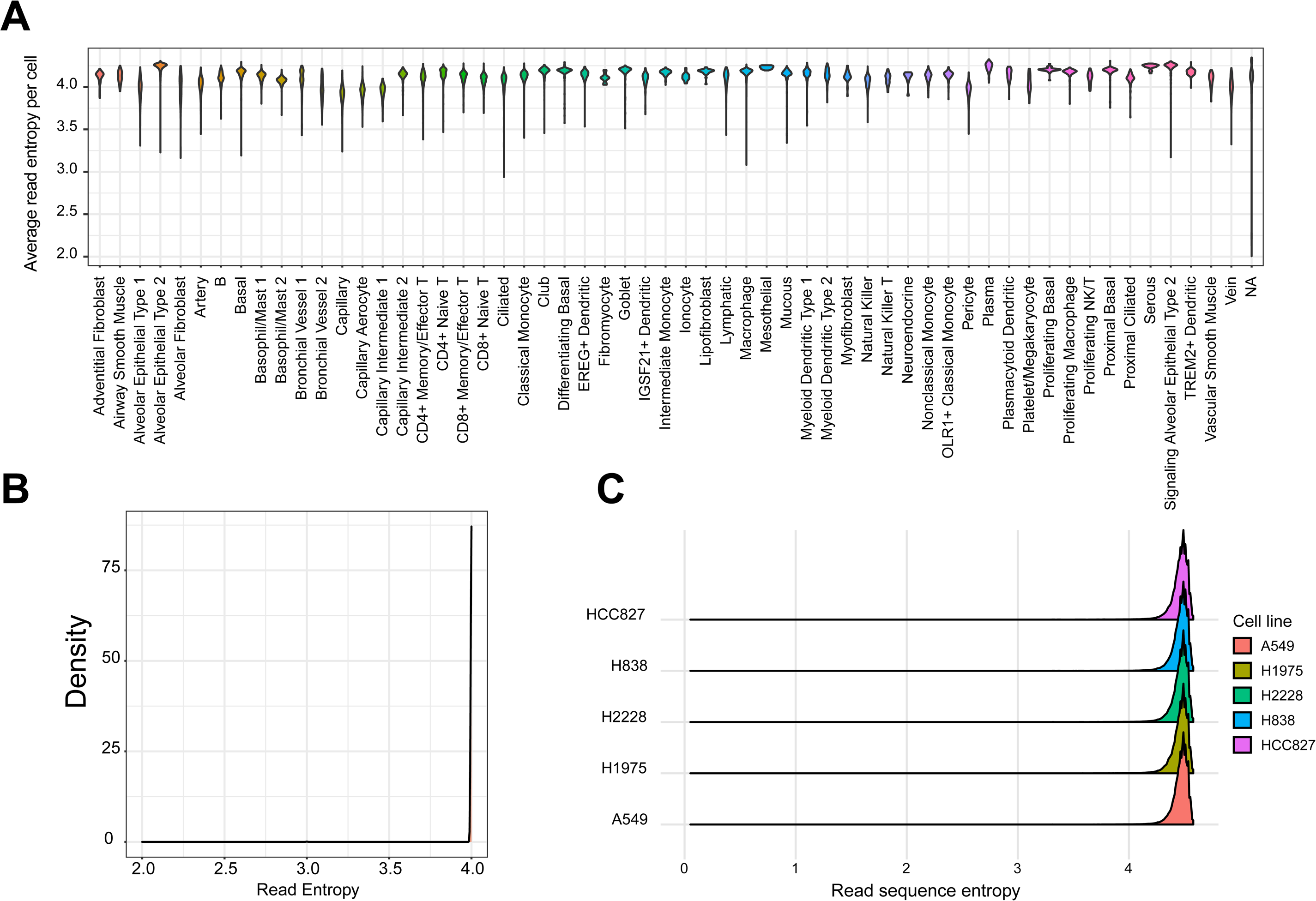
(A) High variation of read sequence entropy in single-cell data. Each violin plot shows the distribution of average read entropy for the cells within a cell type in mouse lemur 10x data. (B) Simulated datasets do not model entropy variation in real data. The density plot shows the read entropy in a simulated dataset ^9^. (C) Low entropy reads in real bulk RNA-seq datasets are less prevalent than single-cell data. The plot shows the read entropy distributions in five cell lines used for generating the benchmarking single-cell dataset ^17^.

**Supplementary Figure 2.**
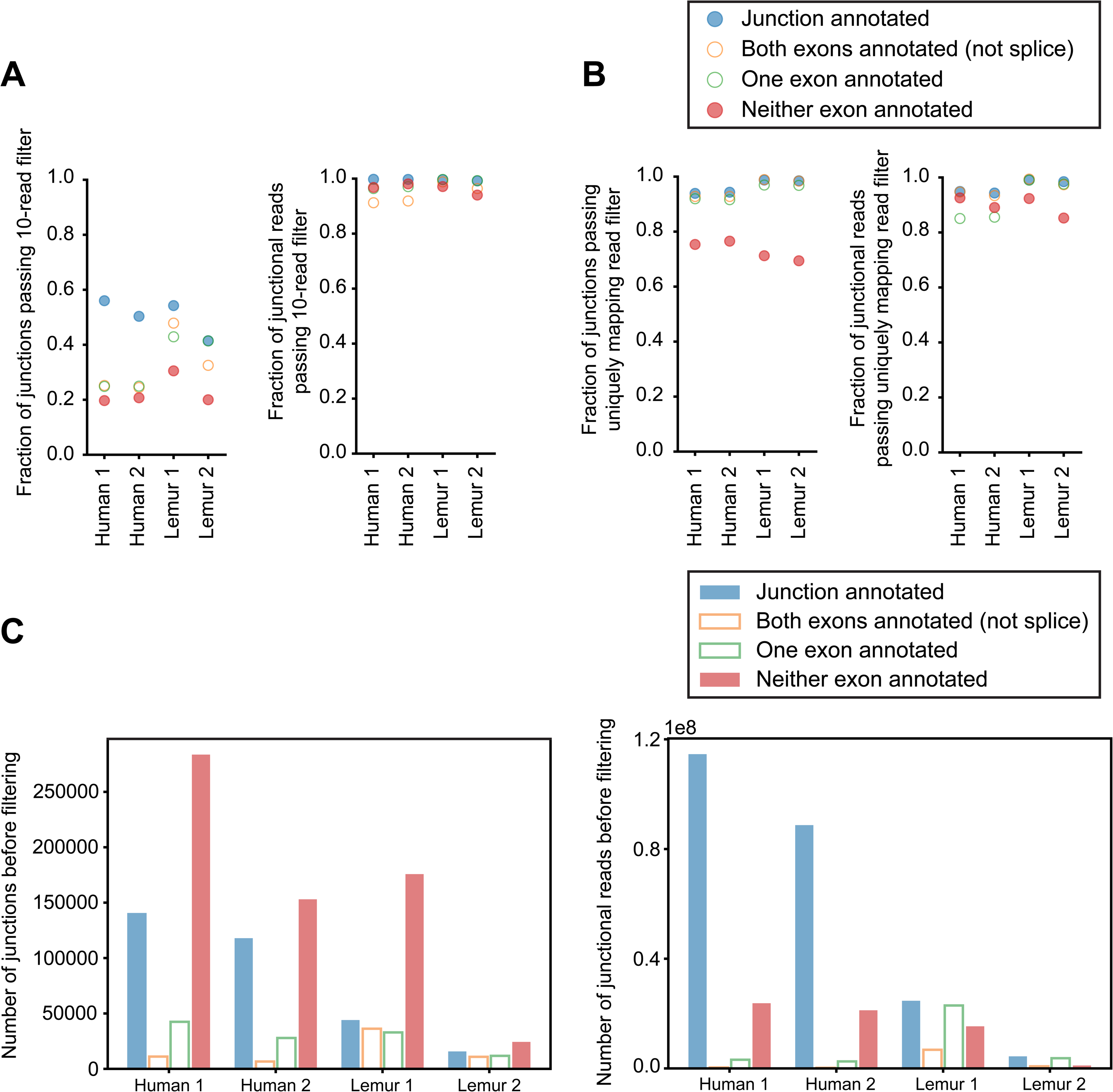
(A) Using a 10 read count threshold causes fewer annotated junctions to be called, and many more unannotated junctional reads to be called [only junctions with at least two reads in the given dataset are plotted]. (B) Including all junctions that have at least one read uniquely mapping to them causes a high fraction of unannotated junctions and unannotated junctional reads to be called [only junctions with at least two reads in the given dataset are plotted]. (C) The number of unannotated junctions is greater than the number of annotated junctions before filtering in all individuals. Lemur individuals have a higher proportion of junctions with one or both exons annotated but without the junction annotated.

**Supplementary Figure 3.**
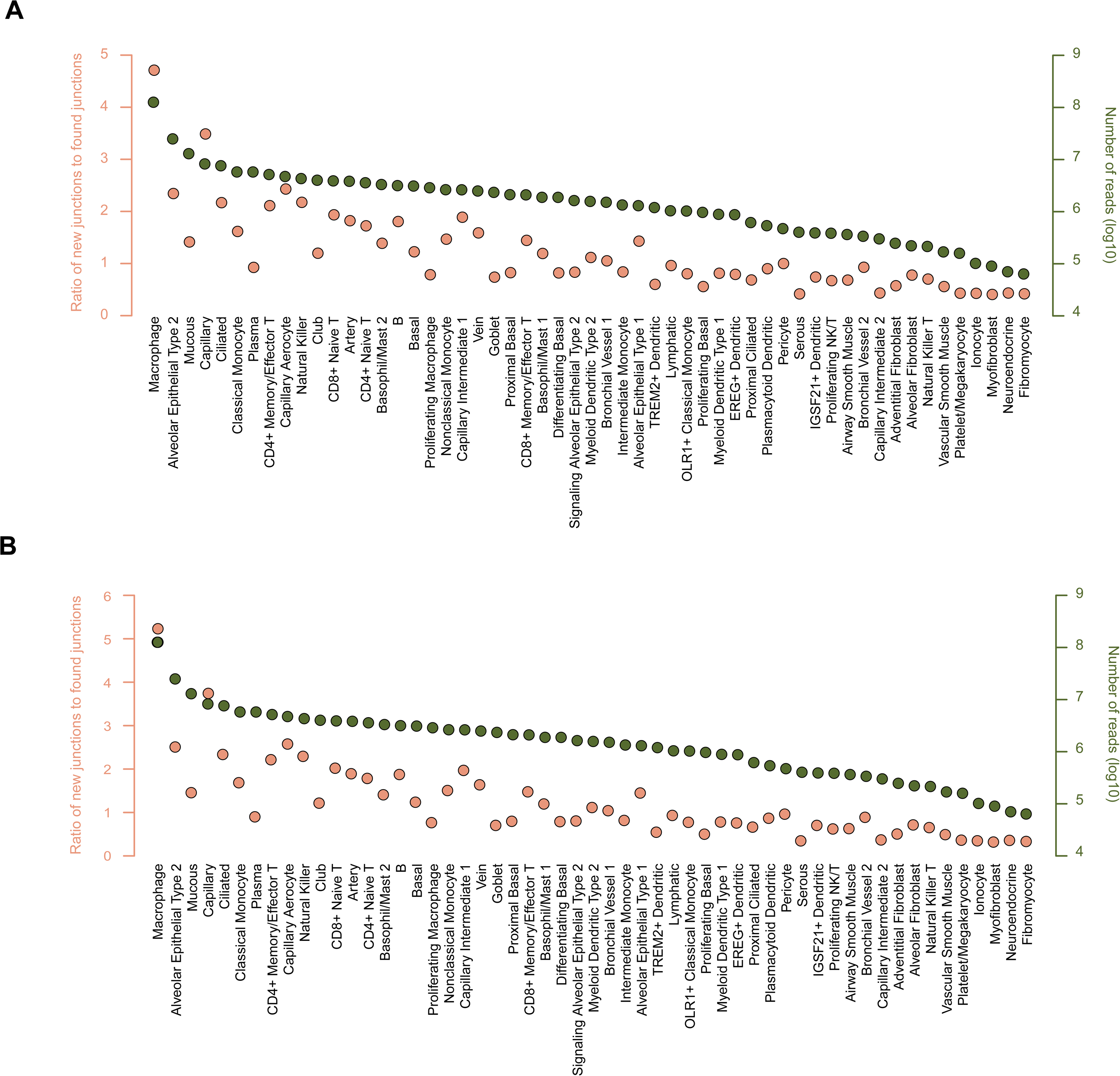
Comparison of the HLCA junctions with two splicing databases: CHESS ^23^ (A) and GTEx ^24^ (B). For each cell type within the HLCA data, the plot shows the ratio of the number of junctions that are not found in the database (defined as new junctions) to the number of junctions that are found in the database (defined as found junctions). The green dots show the number of junctional sequencing reads (in the logarithmic scale) for each cell type.

## List of Supplemental Tables

Table 1 A-G: Breakdown of how many junctions are found in each individual without filtering, after SICILIAN, and after read thresholding.

## Notes

### Competing Interest Statement

The authors have declared no competing interest.

